# Cross-Modal Organization of Facial and Vocal Behavior Revealed by Coordination Metrics Derived from Temporal Alignment

**DOI:** 10.64898/2026.09.26.754599

**Authors:** Yumi Shikauchi, Taiga Naoe, Lara MC Puhlmann, Raffael Kalisch, Fumitaka Homae, Motoaki Nakamura

## Abstract

Facial and vocal behaviors jointly convey affective and communicative information, yet their cross-modal organization remains difficult to characterize quantitatively. We propose a framework for characterizing facial-vocal organization by aligning facial and vocal dynamics using canonical time warping. Multivariate facial action unit trajectories and vocal prosodic trajectories were extracted from recordings of spontaneous speech and aligned in a shared latent space. From the alignment process, we derived three cross-modal coordination metrics: “scaling factor”, reflecting the relative scaling between modalities; “time shift ratio”, reflecting the frequency of temporal shifts; and “time shift magnitude”, reflecting their size. We evaluated these metrics in verbally fluent adults with autism spectrum disorder (ASD; N = 35) and controls (N = 44). The proposed metrics did not differ significantly between diagnostic groups but showed associations with individual differences in communication and anxiety. Scaling factor was linked to trait anxiety in ASD, whereas time shift ratio was associated with communication characteristics across participants. The proposed metrics also showed little association with conventional vocal pitch variability, suggesting that they capture aspects of expressive organization not reflected by vocal variability alone. Our findings support alignment-derived metrics as a complementary approach for characterizing individual variability in facial-vocal expressive behavior.

## I. INTRODUCTION

Communicative behavior is inherently multimodal, with information conveyed through facial and vocal channels that unfold in relation to each other over time. Such cross-modal integration is not limited to phonemic perception, but extends to the interpretation of affective states and communicative intent. Observers draw on facial expressions, vocal prosody, and the temporal relationships between them to infer the emotional states and intentions of others [1,2]. Importantly, this organization is not merely additive; the coordination among expressive channels may itself constitute a behaviorally meaningful signal. Indeed, facial and vocal expressions unfold through different modalities, but their speed and timing do not necessarily coincide. Observers are sensitive to such temporal relationships when integrating audiovisual communicative signals [3]. From a biological perspective, facial and vocal expressions are generated by partially independent but neurally coupled motor systems. In particular, ventral vagal pathways involved in the social engagement system have been proposed to influence both facial musculature and vocal production, suggesting that individual differences in autonomic regulation may be reflected both within each expressive channel and in their temporal relationships [4,5].

A fundamental challenge in understanding multimodal communicative behavior is that expressive organization varies substantially across individuals. Rather than representing noise, such variability may reflect meaningful differences in how communicative information is organized and expressed across facial and vocal channels. Autism spectrum disorder (ASD) provides an informative framework for examining variation in multimodal expressive behavior. Although social communication differences are a defining characteristic of ASD, growing evidence suggests that these differences may be reflected during reciprocal interaction and in facial and vocal expressive behavior [6,7]. However, substantial heterogeneity exists within the autism spectrum, highlighting the importance of considering individual variation beyond diagnostic-group differences. More broadly, expressive behavior can be modulated by individual factors that are not specific to ASD. One such factor is anxiety, which is highly prevalent in individuals with ASD but also manifests in the broader population [8,9]. Elevated anxiety has been associated with alterations in facial expressivity and vocal prosody, suggesting that anxiety may shape multimodal expressive behavior. Together, these observations raise the possibility that variation in expressive behavior lies both within individual modalities and in how facial and vocal expressions are coordinated across modalities. Such cross-modal organization may relate to individual differences in social communication and psychological characteristics.

Recent advances in artificial intelligence have greatly expanded multimodal approaches for affective computing and autism research [10–12]. These approaches have been highly effective for prediction, but typically represent multimodal behavior through modality-specific features, feature fusion, or learned representations. Such representations do not necessarily provide measures that can be directly interpreted regarding how expressive channels relate to one another. Despite the potential importance of temporal relationships between facial and vocal expressions in multimodal communication, few quantitative methods directly characterize how these channels unfold relationally over time [13]. Consequently, interpretable measures of cross-modal expressive organization—measures that describe specific properties of the relationship between facial and vocal expressions—remain underdeveloped. This limits our ability to examine how individual differences in cross-modal expressive organization relate to communicative and affective characteristics.

To address this gap, we propose a framework for characterizing cross-modal expressive organization based on canonical time warping (CTW) [14]. CTW was applied to multivariate facial action unit (AU) trajectories and vocal prosodic feature trajectories extracted from video recordings of spontaneous speech, enabling alignment of expressive dynamics across modalities [15,16] (Fig. 1). As a complementary analysis, we also compared these metrics with vocal pitch variability, which has been widely studied as an acoustic characteristic of speech in ASD [17]. This comparison allowed us to examine whether the proposed cross-modal metrics capture information beyond conventional modality-specific measures. Rather than treating this alignment solely as a preprocessing step, we derived interpretable behavioral descriptors from the alignment process itself, including a scaling factor and metrics quantifying the frequency and magnitude of temporal shifts between modalities. We interpret these measures as complementary indices of cross-modal expressive organization. We evaluated this framework by testing verbally fluent adults with ASD (N = 35) and typically developed controls (TDC) (N = 44). Specifically, we examined whether the proposed coordination metrics were associated with communicative characteristics, assessed using the Communication algorithm score of the Autism Diagnostic Observation Schedule, Second Edition (ADOS-2) [18,19], and with trait anxiety. Rather than focusing solely on diagnostic group differences, we sought to determine whether the proposed coordination metrics capture individual differences in communicative and psychological characteristics.

**Fig. 1.**
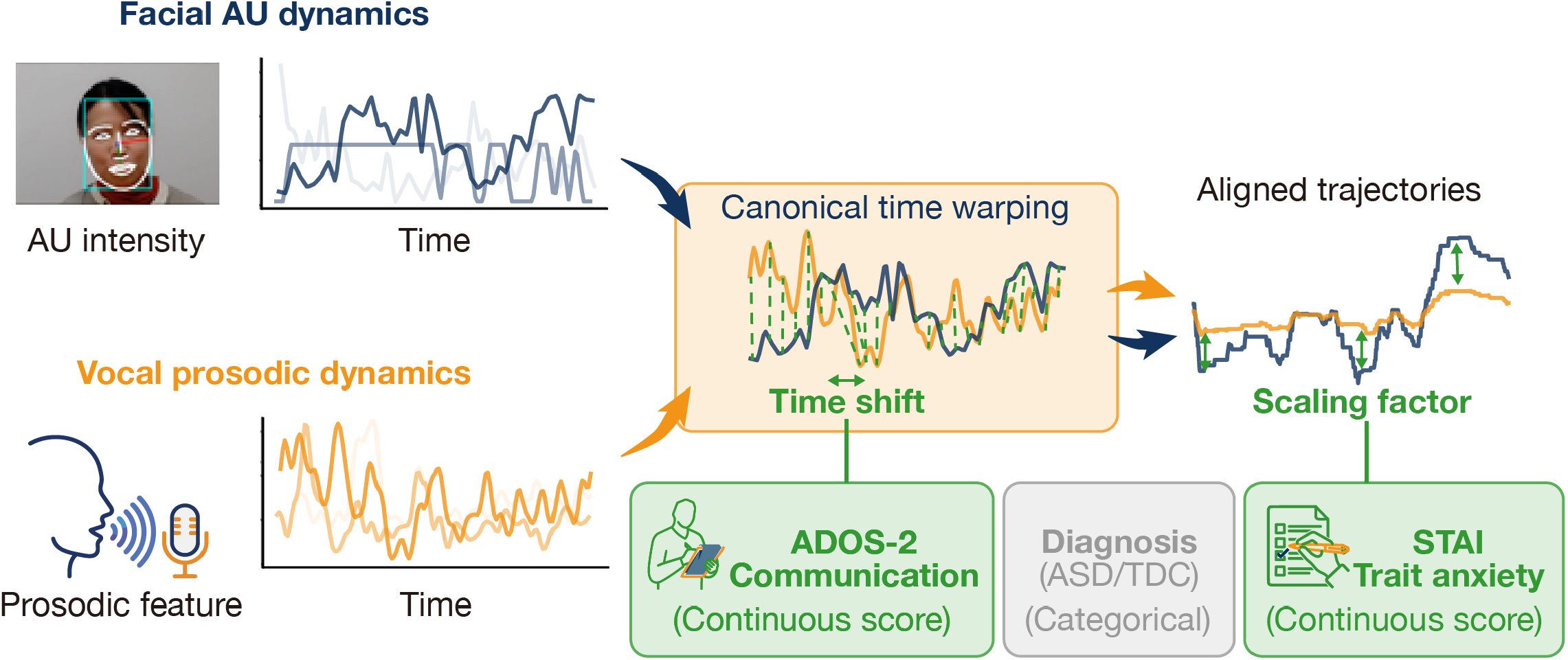
Overview of the study and main findings. Facial action unit (AU) dynamics and vocal prosodic dynamics were extracted from video and audio recordings of spontaneous speech. Canonical time warping (CTW) was applied to align the multivariate facial and vocal trajectories within a shared latent space while compensating for temporal misalignment between modalities. Two classes of coordination metrics were derived from the CTW output: “time shift” metrics, which quantify how frequently and to what extent facial and vocal trajectories are temporally shifted relative to each other, and “scaling factor”, which quantifies the relative magnitude of the two latent trajectories after temporal alignment and correlation maximization. These metrics were subsequently examined in relation to diagnostic group, continuous clinical characteristics, and vocal pitch variability to characterize individual differences in multimodal expressive organization. The illustrative image representing speech was created using the “Generate Vectors” generative AI feature in Adobe Illustrator. ASD: autism spectrum disorder; TDC: typically developed control; ADOS-2: Autism Diagnostic Observation Schedule, Second Edition; STAI: State-Trait Anxiety Inventory.

## II. MATERIALS AND METHODS

### A. Participants

Forty-two individuals with ASD (30 males and 12 females; age: 37 years [IQR: 28–44]) and 52 individuals with TDC (40 males and 12 females; age: 37 years [25–47]) participated in this study. All the participants provided written informed consent. The institutional review board of Showa Medical University approved the recruitment procedures and experimental protocols (approval number: 2023-181-A), which followed the Declaration of Helsinki.

Participants in the ASD group were recruited from Showa Medical University Karasuyama Hospital. Diagnoses of neurodevelopmental disorders were established according to the Diagnostic and Statistical Manual of Mental Disorders, Fifth Edition (DSM-5), based on comprehensive clinical evaluations, including developmental history, current symptoms, life history, and family history. ADOS-2 Module 4 was used as part of the diagnostic assessment. Inclusion criteria for the ASD group were the ability to provide written informed consent and the absence of severe psychiatric, neurological, or physical comorbidities. TDC participants were recruited through public advertisements on a website (www.jikken-baito.com) and through a participant recruitment service provided by SOUKEN Co., Ltd. They were included if they reported no clinically significant psychiatric, neurological, or physical conditions.

Of the 94 participants initially enrolled, 35 individuals with ASD and 44 TDC participants met the eligibility criteria for the present analyses. Participants were excluded because complete ADOS-2 data were unavailable (n = 2), video recordings were unavailable or of poor quality (n = 5), consent was withdrawn (n = 1), the assessment was cancelled (n = 3), the full-scale intelligence quotient (FSIQ) was below 80 (n = 2), no State-Trait Anxiety Inventory (STAI) data were available (n = 1), or, in the TDC group, the ADOS-2 classification was consistent with autism (n = 1). All included participants were video recorded during the storytelling task. The recordings were made at Showa Medical University Karasuyama Hospital between April 2024 and February 2026. Standardized assessments, including ADOS-2 Module 4, Wechsler Adult Intelligence Scale, Fourth Edition (WAIS-IV) or Third Edition (WAIS-III), STAI, and Japanese Adult Reading Test (JART), were obtained either from clinical records or assessments administered as part of this study. Because WAIS scores were not always available (e.g., when prior assessments conducted at other institutions were not accessible), JART was administered to all participants to estimate intellectual functioning. All included participants had JART-estimated IQ scores above 85. Sex (male/female) was self-reported by the participants.

### B. Speech Video Measurements

Participants were seated comfortably in a chair and instructed to relax while watching a silent version of ‘Partly Cloudy,’ a 5.6-min animated movie [20,21]. Before viewing, participants were informed that they would later be asked to describe the movie content, which was presented on a monitor positioned directly in front of the participants.

After viewing, the participants were prompted to describe the content accurately with as much detail as they could. There were no time limits for the storytelling task, allowing participants to speak at their own pace. During this task, facial expressions and speech were recorded using a webcam (Microsoft LifeCam Studio, Microsoft) mounted above the monitor. Video was recorded at 10 fps, and audio was sampled at 44.1 kHz.

### C. ADOS-2 Assessment

To assess individual differences in expressive characteristics, we used the Communication algorithm score, based on DSM-IV, from ADOS-2 Module 4. ADOS-2 is a standardized, semi-structured assessment widely used to diagnose ASD that must be administered by trained and certified professionals [18]. Module 4 is designed for verbally fluent adolescents and adults. The Communication algorithm score was selected because it primarily reflects observable expressive behaviors rather than reciprocal social interaction. All participants completed ADOS-2 Module 4, which was administered by a qualified ADOS-2 examiner of whom there were two.

### D. Preprocessing and Feature Extraction

The following steps were undertaken as preprocessing to prepare multimodal time-series data for subsequent cross-modal alignment analysis. All preprocessing and analyses were performed in Python 3.10.20 using custom scripts and the open-source tools described below. Fig. 2. illustrates the overall pipeline.

**Fig. 2.**
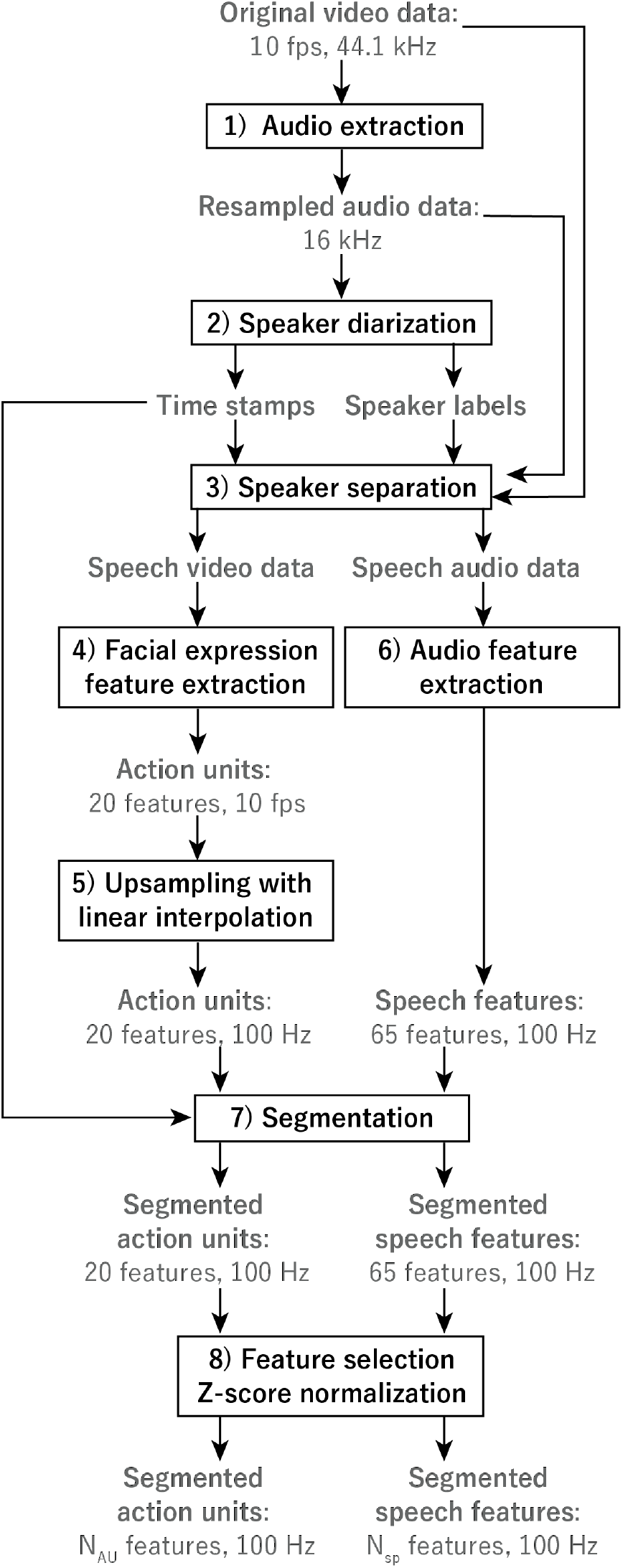
Preprocessing pipeline for multimodal feature extraction and segmentation. Original video recordings (10 fps, 44.1 kHz audio) were decomposed into audio and visual streams. Speech segments were defined using time-stamps and speaker labels, and only segments corresponding to participant speech (i.e., excluding experimenter instructions before and after the task) were retained for analysis. Facial action units (20 features) and acoustic features (65 features) were computed from the video and audio streams, respectively. To enable cross-modal temporal alignment, the facial features were resampled to match the 100-Hz temporal resolution of acoustic features, and both feature sets were segmented into corresponding speech intervals. The extracted features were subsequently screened to remove uninformative dimensions (e.g., low variance, high kurtosis, and monotonic trends), resulting in N_AU_ and N_sp_ features for facial and speech modalities, respectively. These synchronized multimodal time series were used for subsequent CTW-based alignment analysis.

First, speech segments were identified from the video recordings using speaker diarization and separation implemented in OpenWillis v.3.0.5 [22]. Only time periods during which the participant was actively speaking were retained for subsequent analysis. Although the recordings included interactions with an experimenter (e.g., task instructions before and after the speech), only participant-produced speech segments were extracted based on speaker labels to focus on spontaneous solo expression. Based on these segments, we calculated the number of speech segments and the average duration of continuous speech (segment duration) [23,24].

Next, multimodal features were extracted from the speech segments. Facial expression features were obtained from video frames using Py-Feat with its default AU detection model (XGB) [25], yielding 20-dimensional AU representations reflecting facial muscle activity [16]. Acoustic features were extracted from the corresponding audio segments using OpenSMILE with the ComParE_2016 configuration [26], resulting in 65-dimensional feature vectors capturing vocal properties such as intensity, fundamental frequency (F0), and spectral characteristics. Acoustic features were computed using a 60-ms analysis window with a 10-ms frame step, yielding a temporal resolution of 100 Hz. The extracted features were subsequently screened to remove uninformative dimensions based on small variance (< 1 × 10^-6^), high kurtosis (> 10), and monotonic trends.

The resulting 16 to 20 AU features and 23 to 60 speech features were retained for further analysis. Finally, all features were z-normalized within each segment. Following Newby et al. [27], 11 of the 20 AUs (AU09, AU11, AU12, AU14, AU15, AU17, AU20, AU23, AU24, AU25, and AU26) were classified as speech-related based on their designation as AUs involved in speech production.

### E. CTW Analysis

CTW was employed to quantify the correspondence between segmented facial AUs and segmented speech features. CTW combines dynamic time warping (DTW), which aligns a temporally misaligned but correspondingly structured time series, and canonical correlation analysis (CCA), which projects two temporally aligned but unpaired multivariate time series onto a shared latent space [14]. By recursively applying DTW and CCA, CTW enables the alignment of multivariate time series that are temporally misaligned and lack direct correspondence across dimensions. In this study, the dimensionality of the latent space was fixed at 2. This choice was made to ensure stable cross-modal correspondence, as lower-dimensional representations resulted in insufficient alignment, whereas higher-dimensional representations tended to reduce the need for temporal warping. Five iterations of the DTW–CCA optimization were performed. To ensure the numerical stability of CTW, we included only continuous speech segments of at least 800 ms, corresponding to at least eight facial image frames acquired at 10 fps, thereby providing sufficient temporal information for reliable cross-modal alignment. To quantify cross-modal expressive coordination, we extracted three complementary metrics from the CTW output. We estimated a cross-modal scaling factor, defined as the single multiplicative factor that optimally transformed the vocal latent trajectory to match the facial latent trajectory within each segment. A scaling factor of 1 indicates equal amplitudes of facial and vocal latent trajectories. Values > 1 indicate that the vocal trajectory exhibited a smaller amplitude than the facial trajectory, whereas values between 0 and 1 indicate that the vocal trajectory exhibited a larger amplitude. Thus, the scaling factor characterizes the relative gain of facial and vocal expressions after temporal alignment. To characterize temporal asynchrony, we additionally quantified the time shift ratio, defined as the proportion of segments exhibiting non-zero temporal shifts, and the time shift magnitude, defined as the median absolute temporal shift among shifted segments. Together, these measures capture distinct aspects of cross-modal expressive organization, reflecting the relative scaling of facial and vocal latent trajectories as well as the frequency and magnitude of temporal misalignment.

Because facial movements associated with speech production may be mechanically coupled with vocal output, we additionally examined the contribution of speech-related AUs to the CTW latent dimensions to estimate the extent to which cross-modal alignment reflected speech-related facial movements. For each segment and latent dimension, squared face-side CTW-projection weights were normalized across the retained AUs, and the normalized weights of the retained speech-related AUs were summed. Participant-level values were calculated as the median across segments.

### F. Pitch Analysis

To examine whether the proposed cross-modal coordination metrics capture information distinct from a conventional modality-specific measure, we additionally compared them with vocal pitch measures, a commonly used acoustic measure of vocal expression [17]. F0 was estimated for each voiced frame using the OpenSMILE ComParE_2016 feature extraction pipeline. Frames without valid F0 estimates were excluded from the analysis.

Participant-level vocal measures were calculated from all valid F0 estimates pooled across speech segments. Mean pitch was defined as the average F0 across all voiced frames, whereas pitch variability was defined as the standard deviation (SD) of F0 across all voiced frames. To evaluate the robustness of these participant-level measures, we additionally computed pitch statistics by first calculating the mean and SD of F0 within each speech segment and then averaging these values across segments. These alternative measures were highly correlated with the pooled estimates for both mean pitch (Spearman’s *ρ* = 0.986, *p* < 0.01) and pitch variability (*ρ* = 0.740, *p* < 0.01), indicating that participant-level rankings were largely preserved across the two calculation methods.

### G. Statistical Analysis

Group differences in coordination and pitch metrics were assessed using Mann–Whitney U tests. Associations between coordination metrics and clinical measures were evaluated using partial Spearman correlations. Analyses were conducted separately in the ASD and TDC groups and in the combined sample to assess whether observed associations showed a consistent pattern across diagnostic groups. Age, sex, and IQ were included as covariates. FSIQ from the WAIS-IV or WAIS-III was used when available; otherwise, estimated IQ from JART was substituted. Clinical and psychological measures included the ADOS-2 Communication algorithm score and the STAI scores. Analyses involving ADOS-2 Communication were specified a priori based on the study hypothesis that cross-modal expressive coordination would relate to clinically evaluated communicative characteristics. Analyses involving STAI were conducted to examine whether coordination metrics were associated with individual differences differences in anxiety, a factor known to influence vocal and nonverbal expressive behavior. To control for multiple comparisons, *p*-values were adjusted using the Benjamini– Hochberg false discovery rate (FDR) procedure within each family of related tests.

### H. Correlation Structure Analysis

To examine whether the CTW-derived coordination metrics from captured information are distinct from conventional vocal variation, we assessed partial Spearman correlations among pitch variability, scaling factor, time shift ratio, and time shift magnitude, adjusting for age, IQ, sex, and diagnosis. Multiple comparisons across the six pairwise correlations were controlled using the Benjamini–Hochberg FDR procedure.

## III. RESULTS

## A. Participant and Data Characteristics

We first examined demographic, cognitive, and clinical characteristics of the participants (Table I). There were no significant group differences in age, sex, or FSIQ. As expected, ASD participants showed higher ADOS-2 and STAI scores compared to TDC, with higher ADOS-2 Communication algorithm scores indicating greater communication difficulties STAI scores indicating greater trait and state anxiety.

**Table I.** PARTICIPANT CHARACTERISTICS.

| Category | Measure | ASD (n = 35) | TDC (n = 44) | p-value |
| --- | --- | --- | --- | --- |
| Demographic characteristics | Age (years) | 37.000 [27.500–43.500] | 37.000 [25.000–47.000] | 0.859 ($r = 0.021$) |
| | Sex, male/female | 25/10 | 34/10 | 0.739 ($\chi^2 = 0.111$) |
| Clinical and cognitive measures | WAIS-IV FSIQ (n = 25/44) | 111.000 [103.000–120.000] | 116.000 [106.750–130.750] | 0.129 ($r = 0.183$) |
|  | WAIS-III FSIQ (n = 4/0) | 97.000 [91.250–103.000] | N/A | N/A |
| | JART estimated IQ | 113.737 [107.674–117.779] | 111.716 [105.653–117.779] | 0.785 ($r = 0.031$) |
| | ADOS-2 Communication | 3 [2–4] | 1 [0–1] | < 0.010 ($r = 0.772$) |
| | STAI Trait anxiety | 53 [43–63] | 39 [33–46] | < 0.010 ($r = 0.473$) |
| | STAI State anxiety | 39 [32–47] | 35 [31–41] | 0.043 ($r = 0.228$) |

Participant characteristics include demographic and cognitive measures. Values are presented as median and interquartile range (IQR). Group differences were assessed using the Mann–Whitney U test for continuous variables and the chi-square test for categorical variables. Intellectual functioning was confirmed based on available Wechsler Adult Intelligence Scale (WAIS-IV or WAIS-III) scores or Japanese Adult Reading Test (JART) estimates. Significant associations are shown in bold. ASD: autism spectrum disorder; TDC: typically developed controls; FSIQ: full-scale intelligence quotient; IQ: intelligence quotient; ADOS-2: Autism Diagnostic Observation Schedule, Second Edition; STAI: State–Trait Anxiety Inventory.

We next examined segment characteristics and the quality of CTW alignment to evaluate the comparability of the extracted data between groups (Table II). The number of segments per participant did not differ significantly between groups. Segment duration showed a statistically significant difference between groups. However, the effect size was small (*r* = 0.057). After CTW, correlations between cross-modal features were comparable between groups in both latent dimensions, indicating similar alignment quality across groups. We additionally examined the contribution of speech-related facial movements to the CTW latent representations. Speech-related AUs accounted for approximately half of the face-side CTW projection weights in both latent dimensions, with similar contributions across diagnostic groups. This suggests that speech-related facial movements did not differentially influence the CTW-derived representation between groups.

**Table II.** SEGMENT AND ALIGNMENT CHARACTERISTICS.

| Category | Measure | ASD (*n* = 35) | TDC (*n* = 44) | *p*-value |
| --- | --- | --- | --- | --- |
| Segment characteristics | Total number of segments | 866 | 1239 | N/A |
|  | Segments per participant | 15 [6–28] | 21 [5–44] | 0.421 (*r* = 0.091) |
|  | **Segment duration (s)** | **2.455 [1.395–4.297]** | **2.130 [1.300–3.915]** | **0.018 (*r* = 0.051)** |
| Alignment quality | Cross-modal correlation (latent dimension 1) | 0.962 [0.949–0.970] | 0.962 [0.949–0.970] | 0.832 (*r* = 0.024) |
|  | Cross-modal correlation (latent dimension 2) | 0.946 [0.928–0.957] | 0.948 [0.929–0.954] | 0.632 (*r* = 0.054) |
| AU contribution | Speech-related AU contribution (latent dimension 1) | 0.549 [0.512–0.597] | 0.537 [0.503–0.585] | 0.487 (*r* = 0.079) |
|  | Speech-related AU contribution (latent dimension 2) | 0.545 [0.502–0.620] | 0.529 [0.490–0.569] | 0.111 (*r* = 0.180) |

Segment characteristics and quality of canonical time warping (CTW) are summarized for each group. Values are presented as median and interquartile range (IQR). Group differences were assessed using the Mann–Whitney U test. Significant associations are shown in bold. ASD: autism spectrum disorder; TDC: typically developed controls.

### B. Cross-Modal Alignment Metrics

Three participant-level metrics were derived from the CTW alignment: scaling factor, time shift ratio, and time shift magnitude. These metrics characterize complementary aspects of the relative scaling and temporal organization of facial and vocal trajectories.

We compared cross-modal temporal alignment metrics between ASD and TDC participants (Table III). No significant group differences were observed in scaling factor, time shift ratio, or time shift magnitude.

**Table III.** GROUP DIFFERENCES IN CROSS-MODAL TEMPORAL COORDINATION METRICS.

| Metric | ASD (*n* = 35), median [IQR] | TDC (*n* = 44), median [IQR] | *p* (uncorrected) | *p*<sub>FDR</sub> | *r* |
| --- | --- | --- | --- | --- | --- |
| Scaling factor | 1.335 [0.904–1.640] | 1.203 [0.572–1.524] | 0.499 | 0.741 | 0.077 |
| Time shift ratio | 0.889 [0.846–1.000] | 0.938 [0.880–1.000] | 0.401 | 0.741 | 0.093 |
| Time shift magnitude | 12.171 [10.043–14.158] | 11.904 [9.734–13.918] | 0.741 | 0.741 | 0.038 |

The Benjamini–Hochberg false discovery rate (FDR) correction was applied across the three alignment metrics. ASD: autism spectrum disorder; TDC: typically developed control; IQR: interquartile range.

We next examined whether cross-modal coordination metrics were associated with clinically evaluated communicative characteristics and anxiety-related traits. Partial Spearman correlations were calculated while controlling for age, sex, and IQ (Fig. 3 and TABLE IV). Within the ASD group, scaling factor was positively associated with trait anxiety (ρ = 0.418, pFDR = 0.026). Because a scaling factor of 1 indicates balanced facial and vocal amplitudes, whereas values > 1 indicate relatively smaller vocal dynamics, this result suggests that higher trait anxiety was associated with reduced vocal variability relative to facial expression. No significant associations were observed between scaling factor and ADOS-2 Communication scores or state anxiety.

**TABLE IV.**
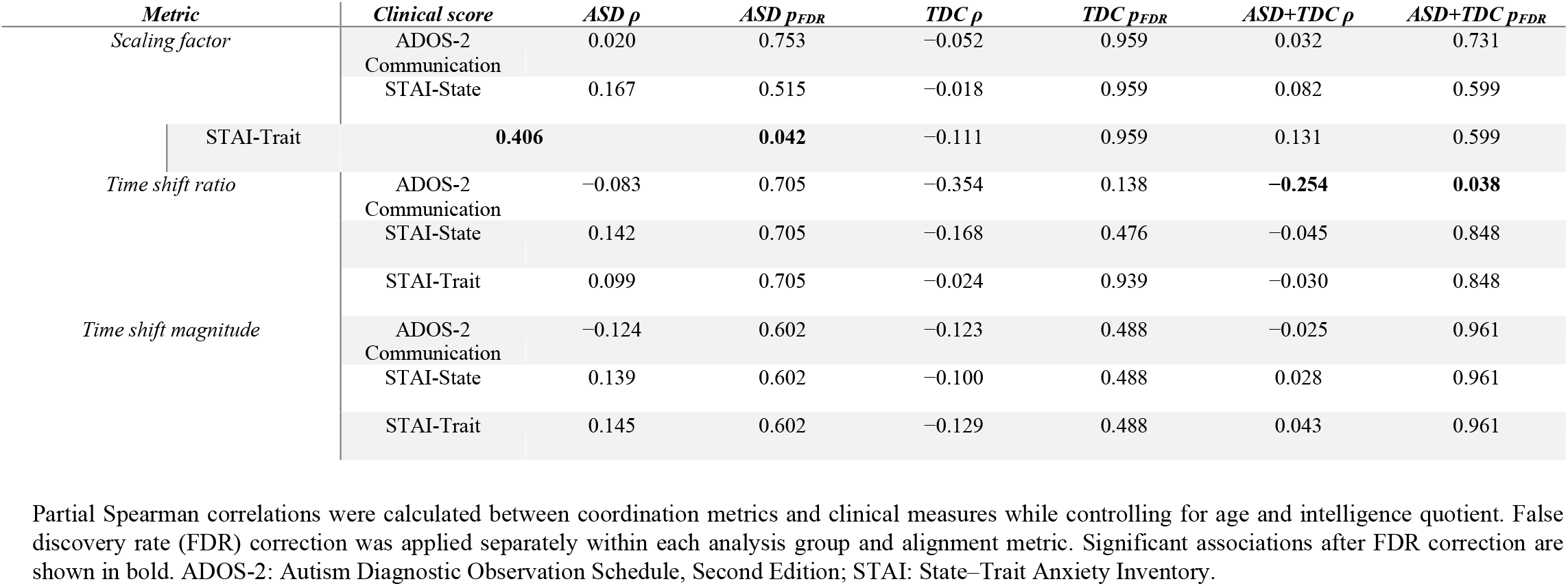
ASSOCIATIONS BETWEEN CROSS-MODAL CORERLATION METRICS AND CLINICAL MEASURES IN CROSS-MODAL.

**Fig. 3.**
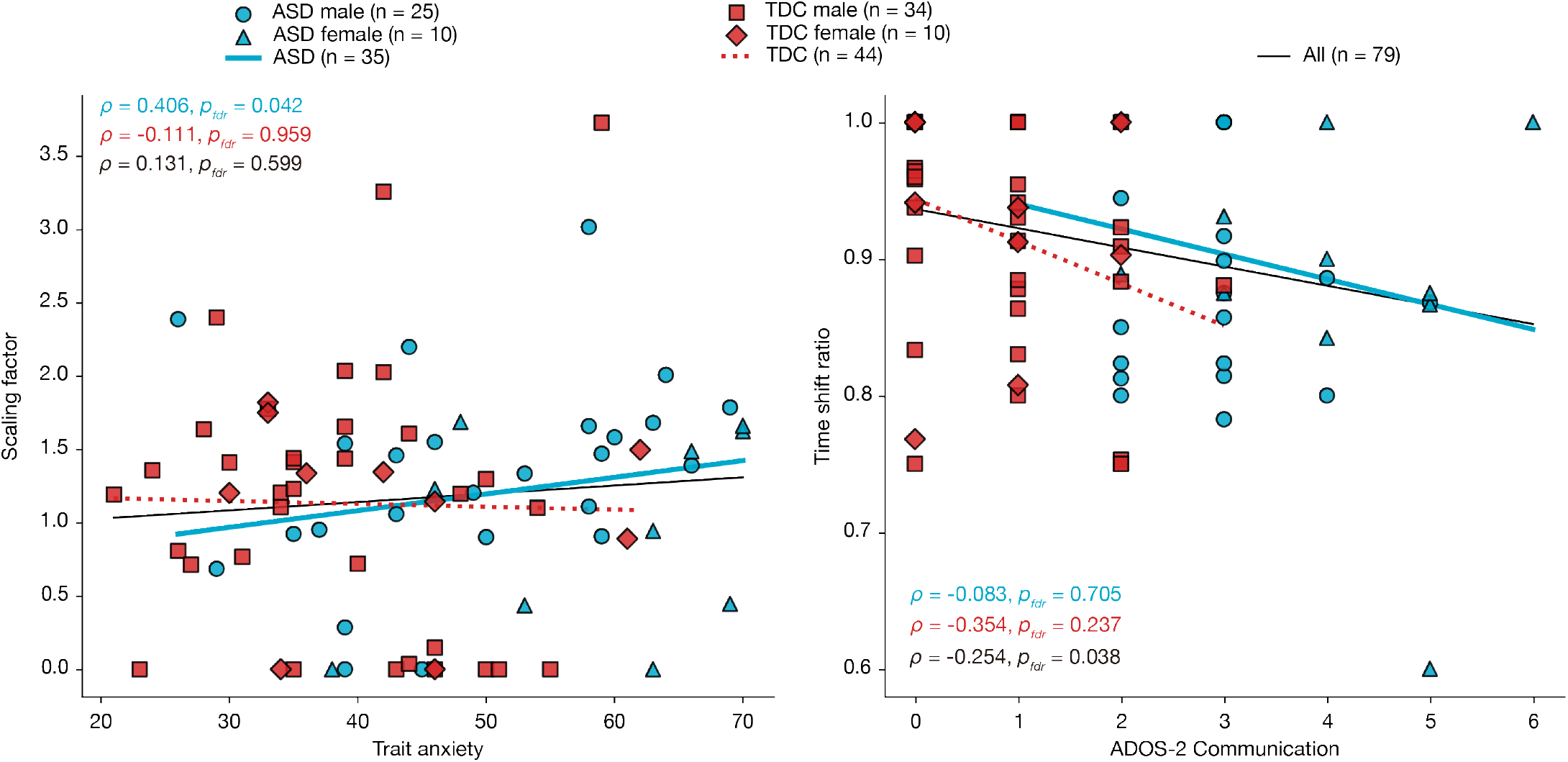
Distinct associations of cross-modal coordination metrics with anxiety and communication. Scatter plots illustrate the significant associations identified by partial Spearman correlation analyses controlling for age, sex, and intelligence quotient. Left, scaling factor as a function of STAI-Trait anxiety score within the ASD group. Right, time shift ratio as a function of ADOS-2 Communication score within the combined group. Each point represents one participant. Solid cyan lines indicate linear trends for ASD participants, dashed red lines indicate linear trends for TDC participants, and the solid black line represents the linear trend across all participants. Trend lines were derived from simple linear regression (Pearson correlation) for visualization only. Correlation statistics are reported in TABLE 4.

Measures reflecting temporal shifts between modalities were associated with communicative characteristics. Within the combined samples, time shift ratio showed a significant negative association with ADOS-2 Communication scores (*ρ* = −0.261, *p*_FDR_ = 0.036), indicating that individuals with greater communication difficulties exhibited fewer temporally shifted facial–vocal segments. The direction of the association was consistent across the two groups.

### C. VOCAL PITCH CHARACTERISTICS

To determine whether the observed associations were specific to cross-modal coordination rather than reflecting variability within a single expressive modality, we performed analogous analyses using conventional vocal pitch measures [16]. Because increased vocal pitch variability has previously been reported in ASD, we first characterized vocal pitch in the present sample. The pitch mean was numerically higher in the ASD group (137.68 [118.12–227.15] Hz) than in the TDC group (132.44 [116.62–166.04] Hz), although this difference was not statistically significant (*p* = 0.487, *r* = 0.079). Likewise, pitch SD was comparable between groups (ASD: 22.47 [17.37– 31.84] Hz, TDC: 21.53 [17.18–32.15] Hz; *p* = 0.972, *r* = 0.004).

Pitch mean was not significantly associated with any clinical measure in the ASD, TDC, or combined samples (Table V). Pitch SD was significantly associated with anxiety within the TDC group (state anxiety: *ρ* = 0.514, *p*_FDR_ < 0.010; trait anxiety: *ρ* = 0.323, *p*_FDR_ = 0.049). These associations were not significant in the ASD or combined samples. There was no significant correlation between pitch metrics and ADOS-2 Communication scores.

**Table V.**
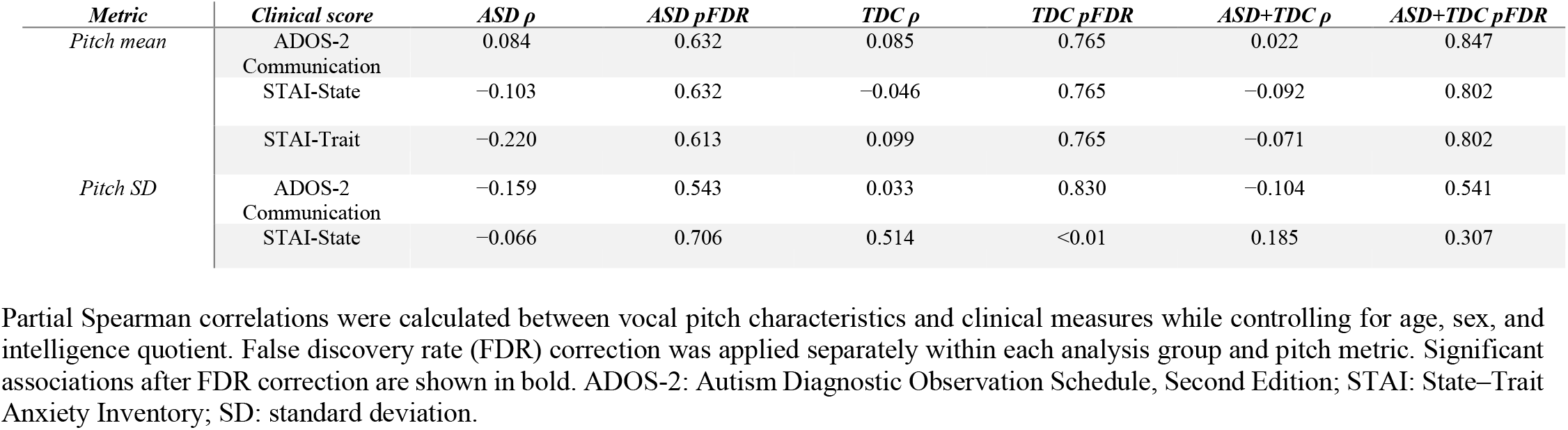
ASSOCIATIONS BETWEEN VOCAL PITCH CHARACTERISTICS AND CLINICAL MEASURES TEMPORAL COORDINATION METRICS.

Because sex differences in vocal pitch metrics are well established, we additionally compared all expressive metrics between males and females within each diagnostic group (Table VI). As expected, both pitch mean and pitch SD differed markedly by sex in both ASD and TDC participants (all *p* < 0.01), whereas no significant sex differences were observed for the proposed cross-modal coordination metrics.

**Table VI.** SEX DIFFERENCES IN CROSS-MODAL COORDINATION METRICS AND VOCAL PITCH CHARACTERISTICS WITHIN EACH DIAGNOSTIC GROUP.

| Group | Metric | Male | Female | $p$ | $r$ |
| --- | --- | --- | --- | --- | --- |
| ASD | Scaling factor | 1.389 [0.923–1.658] (n=25) | 1.086 [0.438–1.588] (n=10) | 0.298 | 0.179 |
| ASD | Time shift ratio | 0.899 [0.824–1.000] (n=25) | 0.882 [0.869–0.923] (n=10) | 0.751 | 0.056 |
| ASD | Time shift magnitude | 12.475 [10.647–16.229] (n=25) | 9.143 [6.507–12.944] (n=10) | 0.065 | 0.315 |
| ASD | Pitch mean | **124.624 [116.7192–138.7879]** (n=25) | **234.6384 [225.9859–245.6475]** (n=10) | **<0.01** | **0.673** |
| ASD | Pitch SD | **13.502 [11.09810–16.0576]** (n=25) | **27.698 [23.3889–29.767]** (n=10) | **<0.01** | **0.605** |
| TDC | Scaling factor | 1.200 [0.289–1.565] (n=34) | 1.269 [0.954–1.458] (n=10) | 0.856 | 0.030 |
| TDC | Time shift ratio | 0.934 [0.879–1.000] (n=34) | 0.939 [0.905–1.000] (n=10) | 0.618 | 0.076 |
| TDC | Time shift magnitude | 12.210 [10.270–14.343] (n=34) | 9.765 [9.585–11.758] (n=10) | 0.101 | 0.249 |
| TDC | Pitch mean | **122.2657 [110.640–131.382]** (n=34) | **241.9465 [216.960–248.9485]** (n=10) | **<0.01** | **0.701** |
| TDC | Pitch SD | **12.264 [9.2657–15.5081]** (n=34) | **33.900 [31.09910–40.4364]** (n=10) | **<0.01** | **0.625** |

Partial Spearman correlations were calculated between vocal pitch characteristics and clinical measures while controlling for age, sex, and intelligence quotient. False discovery rate (FDR) correction was applied separately within each analysis group and pitch metric. Significant associations after FDR correction are shown in bold. ADOS-2: Autism Diagnostic Observation Schedule, Second Edition; STAI: State-Trait Anxiety Inventory; SD: standard deviation.

These findings indicate that vocal pitch variability primarily reflects anxiety-related individual differences, whereas communicative characteristics were associated only with the proposed cross-modal coordination metrics. Together with the absence of significant sex differences in the cross-modal metrics despite the marked sex differences in vocal pitch, these findings suggest that the proposed metrics capture aspects of expressive organization that are not reflected by conventional vocal pitch measures alone.

### D. Interrelationships Among Expressive Metrics

To further characterize the relationships among expressive metrics, we analyzed partial correlations among vocal pitch variability and the three proposed cross-modal coordination metrics across the full sample (Fig. 4). The analyses controlled for age, IQ, sex, and diagnosis. Pitch variability was not significantly associated with scaling factor (*ρ* = 0.040, *p*_FDR_ = 0.735), time shift ratio (*ρ* = −0.086, *p*_FDR_ = 0.554), or time shift magnitude (*ρ* = −0.116, *p*_FDR_ = 0.483). Among the cross-modal coordination metrics, scaling factor was positively associated with time shift ratio (*ρ* = 0.383, *p*_FDR_ = 0.004), whereas the other pairwise associations were not statistically significant. These findings further indicate that the proposed cross-modal coordination metrics capture aspects of expressive organization that are not reflected by conventional vocal pitch variability alone.

**Fig. 4.**
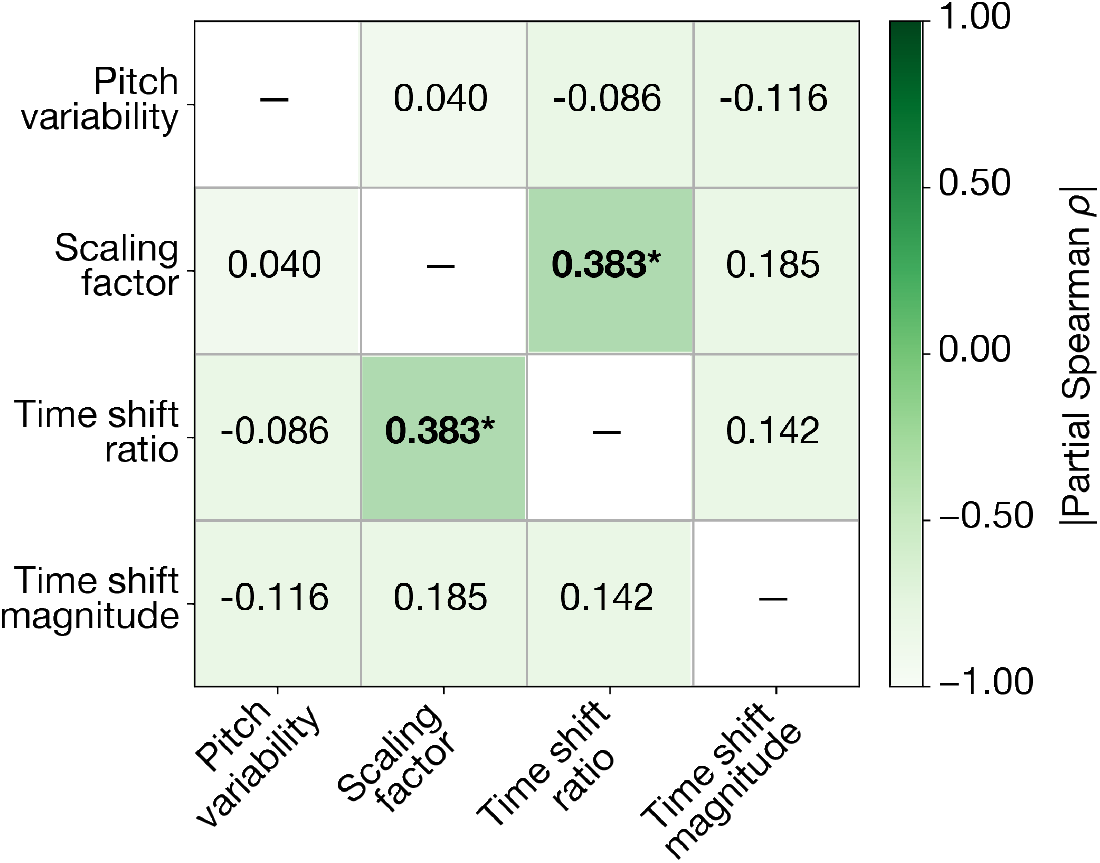
Relationships among cross-modal coordination metrics and vocal pitch variability. Partial Spearman correlations among pitch variability, scaling factor, time shift ratio, and time shift magnitude were calculated across all participants while controlling for age, intelligence quotient, sex, and diagnosis. Values indicate partial Spearman correlation coefficients (*ρ*). An asterisk indicates an association that remained significant after false discovery rate correction across the six pairwise correlations.

## IV. DISCUSSION

We quantified facial–vocal coordination using CTW-derived metrics and examined their relationships with diagnosis and individual communicative and psychological characteristics. Although these metrics did not distinguish ASD from TDC participants, they showed selective associations with clinical and psychological measures. Scaling factor was associated with trait anxiety within the ASD group, whereas time shift ratio was associated with ADOS-2 Communication algorithm scores across participants. By contrast, conventional vocal pitch variability was associated primarily with anxiety within TDC and showed pronounced sex differences. Furthermore, pitch variability showed little association with the proposed cross-modal coordination metrics after accounting for age, IQ, sex, and diagnosis. Together, these findings suggest that the proposed metrics capture aspects of expressive organization not reflected by vocal pitch variability alone and may provide a complementary approach to characterizing individual variability in expressive behavior.

The two coordination metrics associated with clinical measures may reflect different aspects of expressive organization. Higher scaling factors were associated with greater trait anxiety in ASD, suggesting relatively reduced vocal dynamics compared with facial expression, whereas time shift ratio was associated with communicative characteristics. Although temporal shifts may intuitively be interpreted as poorer coordination, spontaneous speech is not produced synchronously across modalities. Speakers continuously formulate linguistic content while simultaneously producing facial and vocal behaviors. This makes modest temporal offsets between modalities a natural feature of expressive communication. Under this interpretation, the association between higher time shift ratios and lower ADOS-2 Communication scores suggests that some degree of temporal flexibility between facial and vocal expressions may be related to more effective communicative behavior. Importantly, ADOS-2 Communication algorithm scores are based on standardized behavioral observations by trained examiners. Although the ADOS-2 does not directly assess cross-modal timing, this association raises the possibility that temporal organization across facial and vocal channels relates to aspects of expressive behavior that are salient in observer-based evaluations of communication. The association was in the same direction in both groups.

### A. Focusing on Individual Expressive Characteristics

A key methodological decision in the present study was to focus on the ADOS-2 Communication algorithm score rather than the revised Social Affect (SA) score [28], based on DSM 5. The Communication algorithm score includes several items that directly reflect observable expressive behaviors, such as intonation, gestures, facial expression, and conversation. In contrast, the SA score integrates these behaviors with broader aspects of reciprocal social interaction. Expressive characteristics reflect how an individual externally manifests internal states such as emotions and intentions. In social contexts, however, these characteristics become intertwined with interactional processes that also depend on the behavior of others. Because the present study aimed to quantify individual characteristics of expressive behavior, we considered the Communication algorithm score a more direct behavioral reference than the broader SA construct.

Additionally, our findings suggest that cross-modal coordination between vocal and facial expressions was associated with individual expressive characteristics during the solo speech task, which involved minimal interactivity. This may represent a practical advantage for the development of digital markers, as expressive behavior can be assessed from recordings produced without an interaction partner. Such recordings are increasingly common in everyday settings, including video messages and personal videos. Whether the proposed metrics remain informative during interactive communication is an important question for future research. Video-mediated communication provides a particularly relevant intermediate setting because it preserves verbal interaction while reducing some of the nonverbal and interactional cues available in face-to-face conversation [29,30]. Evaluating cross-modal coordination in video-mediated communication, and subsequently extending the framework to quantify cross-individual coordination during face-to-face interaction, will help clarify how expressive organization changes across communicative contexts.

Beyond differences in communicative context, expressive behavior is also influenced by the speaker’s psychological state. This may help explain the observed association between pitch variability and state anxiety. In TDC participants, greater pitch variability was associated with higher state anxiety. This finding differs from the negative association reported by Laukka et al. [31], who examined only the initial 10 s of speech during a highly socially evaluative speaking task. Previous work has also shown that pitch variability changes with increasing task demands [32]. Thus, differences in the internal states elicited by different experimental conditions may contribute to variation in vocal expression and may partly account for inconsistencies across studies. In addition, questionnaire-based anxiety may not deterministically map onto the same underlying experiences or behavioral expressions across individuals. Variability in this correspondence may also contribute to the observed relationship between anxiety and vocal expression [33,34]. Future studies should therefore examine both facial and vocal behavior across a broader range of communicative situations, particularly interactive conversations. Incorporating subjective and physiological measures of anxiety could further clarify how internal psychological states are related to natural social expression [35].

### B. Structure of Multimodal Expressive Coordination

The relationships among the expressive metrics provide further insight into what aspects of multimodal organization are captured by the proposed measures. After controlling for age, IQ, sex, and diagnosis, pitch variability showed little association with any of the three cross-modal coordination metrics. This suggests that the proposed metrics capture aspects of expressive organization that are not reflected by conventional vocal pitch variability alone. Among the cross-modal metrics, scaling factor was positively associated with time shift ratio, whereas the other pairwise associations were not significant. Thus, although scaling factor and time shift ratio characterize distinct properties of face–voice coordination, their moderate association suggests that relative scaling and temporal organization are not entirely independent aspects of multimodal expression.

An important question is which aspects of cross-modal organization contribute to effective communication and how their alteration relates to communicative difficulties. Previous studies have suggested that relationships between behavioral modalities may carry clinically relevant information. For example, Sorensen et al. [36] reported weaker facial–vocal coordination in children with ASD during emotional speech, as quantified by Granger causality. At first glance, this may appear inconsistent with our finding that greater communication difficulties were associated with a lower time shift ratio. However, temporal shifts detected by CTW do not necessarily indicate poorer coordination. Rather, they indicate that corresponding facial and vocal dynamics unfold with relative temporal offsets and therefore require temporal adjustment for optimal alignment. Such temporal offsets may be a natural component of multimodal expression rather than a failure of coordination. Thus, the temporal offsets quantified by CTW may reflect a different aspect of face–voice coordination from the dynamic dependencies captured by Granger causality.

Rather than replacing conventional facial or vocal measures, the proposed framework provides a complementary way to characterize how expressive behaviors are organized across modalities, thereby offering an interpretable computational framework for studying individual variability in natural communication.

## ACKNOWLEDGMENTS

We extend our heartfelt thanks to Prof. Yukie Nagai and Prof. Masanori Hariyama for their invaluable contributions and insightful discussions that greatly shaped the initial direction of this research. We would like to express our gratitude to Noriko Ishimura, Mika Kato, and Taku Sato for their contributions to recruiting participants for this study. We are also grateful to Sayuri Takeda and Maiko Inoue for their contributions to the clinical assessments, including administration of the ADOS-2. Finally, we thank Dr. Shohei Tsuchimoto for technical assistance with organizing the analysis code. OpenAI ChatGPT was used for language editing and grammar enhancement of the manuscript. The “Generate Vectors” generative AI feature in Adobe Illustrator was used to create an illustrative image representing speech in Fig. 1. All AI-assisted content was reviewed and approved by the authors.

